# Tumbly: An Open-Source Home-Cage Device for Time-Restricted Feeding in Mice

**DOI:** 10.64898/2026.10.19.760239

**Authors:** Mason R. Barrett, Cynthia Chen, Winston Li, Matt Gaidica, Amy K. Wagner, T. John Wu, Erik S. Musiek, Alexxai V. Kravitz

**Affiliations:** Department of Psychiatry, Washington University in St. Louis, St. Louis, MO, USA; Department of Biomedical Engineering, Washington University in St. Louis, St. Louis, MO, USA; Department of Neurology, Washington University in St. Louis, St. Louis, MO, USA; Department of Gynecologic Surgery and Obstetrics, Uniformed Services University of the Health Sciences, Bethesda, MD, USA; Department of Neuroscience, Washington University in St. Louis, St. Louis, MO, USA; Department of Physical Medicine & Rehabilitation, University of Pittsburgh School of Medicine, Pittsburgh, PA, USA; Department of Neuroscience, University of Pittsburgh School of Arts and Sciences, Pittsburgh, PA, USA; Department of Bioengineering, University of Pittsburgh School of Engineering, Pittsburgh, PA, USA; Safar Center for Resuscitation Research, University of Pittsburgh School of Medicine, Pittsburgh, PA, USA; Department of Anesthesiology, Washington University in St. Louis, St. Louis, MO, USA

## Abstract

Time-restricted feeding (TRF) is a common tool for studying metabolism, circadian rhythms, aging, and neurodegeneration. Here, we describe a new device called Tumbly, an automated device for performing TRF studies in mouse home-cages. Tumbly controls access to rodent diets at user-set times via a programmable servo-driven food hopper. Tumbly is also battery powered and compatible with multiple types of vivarium home-cages, due to a removable faceplate that can be customized to fit different cage dimensions without modification. Finally, Tumbly is open-source and affordable, with all design files available online, allowing researchers to build and customize it. As a validation test, we deployed Tumbly in two labs at separate institutions, one with singly housed mice (n = 6, within-subject) and one with group-housed mice (n = 22, between-group). We measured daily body weight and food intake. In both labs, mice on an 8-hour TRF schedule maintained their body weight and ate similar amounts of food compared to ad libitum feeding. Our quantified error rates were also low, with no device failures over >1000 hours of operation. We conclude that Tumbly provides a reliable, low-cost option for running TRF studies in mice.

**Significance Statement:** Time-restricted feeding (TRF) studies are often run by hand, moving food or animals between cages at fixed times to restrict feeding. This approach limits the study cohort size to what experimenters can manage, requires someone to be present to control food at inconvenient hours, and can add stress to the animals from repeated experimenter interventions. Tumbly removes these limitations by automating the TRF procedure. Tumbly is also open-source and can be modified to adapt it to more complex monitoring experiments, such as adding in-cage temperature or humidity sensors. Finally, because Tumbly is affordable, it makes automation of TRF both more accessible to labs with smaller budgets and scalable for researchers that need to run high throughput TRF studies.

## Introduction

The timing of food intake can both positively and negatively impact health. A landmark study found that restricting high-fat diet access to the inactive phase exacerbated an obesity phenotype in mice, compared to when mice were given the same high-fat diet in their active phase (Arble et al., 2009). This finding raised the intriguing possibility that obesity was controlled not only by what mice were eating, but also by when they were eating it. Restricting feeding to specific hours of the circadian cycle is termed Time-Restricted Feeding (TRF), and the protective effects of TRF on obesity have now been replicated across multiple rodent studies. Restricting high-fat intake to only during an 8 h window in the active phase protected mice from high-fat diet-induced obesity, hyperinsulinemia, hepatic steatosis, and inflammation, importantly without reducing caloric intake (Hatori et al., 2012). Contemporaneous work also demonstrated that restricting high-fat diet to the active phase improved body weight, insulin sensitivity, and circulating metabolic markers (Sherman et al., 2012). TRF has even been shown to drive weight loss and improve metabolic function in mice with pre-existing obesity, expanding its role into reversing obesity, and these benefits held across high-fat, high-fat high-sucrose, and high-fructose diets (Chaix et al., 2014). In humans, results have been more mixed. Early time-restricted eating improved insulin sensitivity in men with prediabetes without weight loss (Sutton et al., 2018), and a 30-day, 8-hour eating window led to weight loss in nearly all participants of a small single-arm trial (Chen et al., 2024). However, time-restricted eating did not produce more weight loss than calorie restriction alone in a randomized trial of adults with obesity (Liu et al., 2022).

In recent years, there has been increased recognition of the beneficial effects of TRF beyond controlling body weight. Restricting feeding to the active phase extended lifespan in mice (Acosta-Rodríguez et al., 2022) and reduced amyloid pathology and improved memory in mouse models of both Alzheimer’s disease (King et al., 2023; Whittaker et al., 2023) and Huntington’s disease (Wang et al., 2018). These preclinical findings have produced excitement for leveraging TRF to modify disease progression in neurodegenerative diseases, both for understanding circadian biology and recently as a promising therapeutic intervention in people with Alzheimer’s disease (Zhao et al., 2025). Clinical evidence in neurodegenerative disease is still limited, but early trials are testing whether time-restricted eating is feasible and beneficial in older adults and people with cognitive impairment (Zhao et al., 2025).

The standard method for applying TRF has been to manually control access to food for animals, by adding and removing food to their cages or transferring animals between cages with and without food at set times of day (Xin et al., 2021; Yan et al., 2024). This intervention is time-consuming for researchers, requiring them to intervene at specific hours to enforce the TRF schedule. In addition, daily experimenter intervention and caging changes can introduce stress to mice (Rasmussen et al., 2011). Automated systems can reduce the handling stress, but can be expensive and are often designed to be compatible with specific caging (Acosta-Rodríguez et al., 2017; Tilston et al., 2019). Several open-source solutions have also been designed to automate TRF experimentation, and to allow users more flexibility (Sahasrabudhe et al., 2022; Farré et al., 2024; Lê et al., 2025). Tumbly attempts to complement existing systems in several ways. Most notably, Tumbly is battery powered so it can live in the home cage without external power. This enables Tumbly to run effectively using rack-mounted vivarium caging without modification, including individually ventilated caging as we use in the present study. We also designed Tumbly to have a removable “faceplate” to allow it to be compatible with multiple types of caging after modifying only the 2D faceplate. Finally, Tumbly is affordable and relatively easy to build without specialized tools, allowing any lab to build it to perform automated TRF experiments.

## Materials and Methods

### Device overview

Tumbly is an open-source home cage TRF device that uses a programmable hopper to control access to food on a user-programmed schedule (Fig. 1A; Movie 1). The 3D printed body was designed to be small enough to fit inside most rodent home-cages (Fig. 1B), while a customizable “faceplate” snaps onto the body to accommodate different home-cage internal sizes (Fig. 1C, D). Tumbly’s printed parts are made from PLA, a common low-toxicity plastic, and the electronics are enclosed so mice cannot reach them. The device is assembled with 3D printed parts and hardware, as detailed in Table 1 and https://github.com/KravitzLabDevices/Tumbly/wiki/Parts-List.

**Figure 1.**
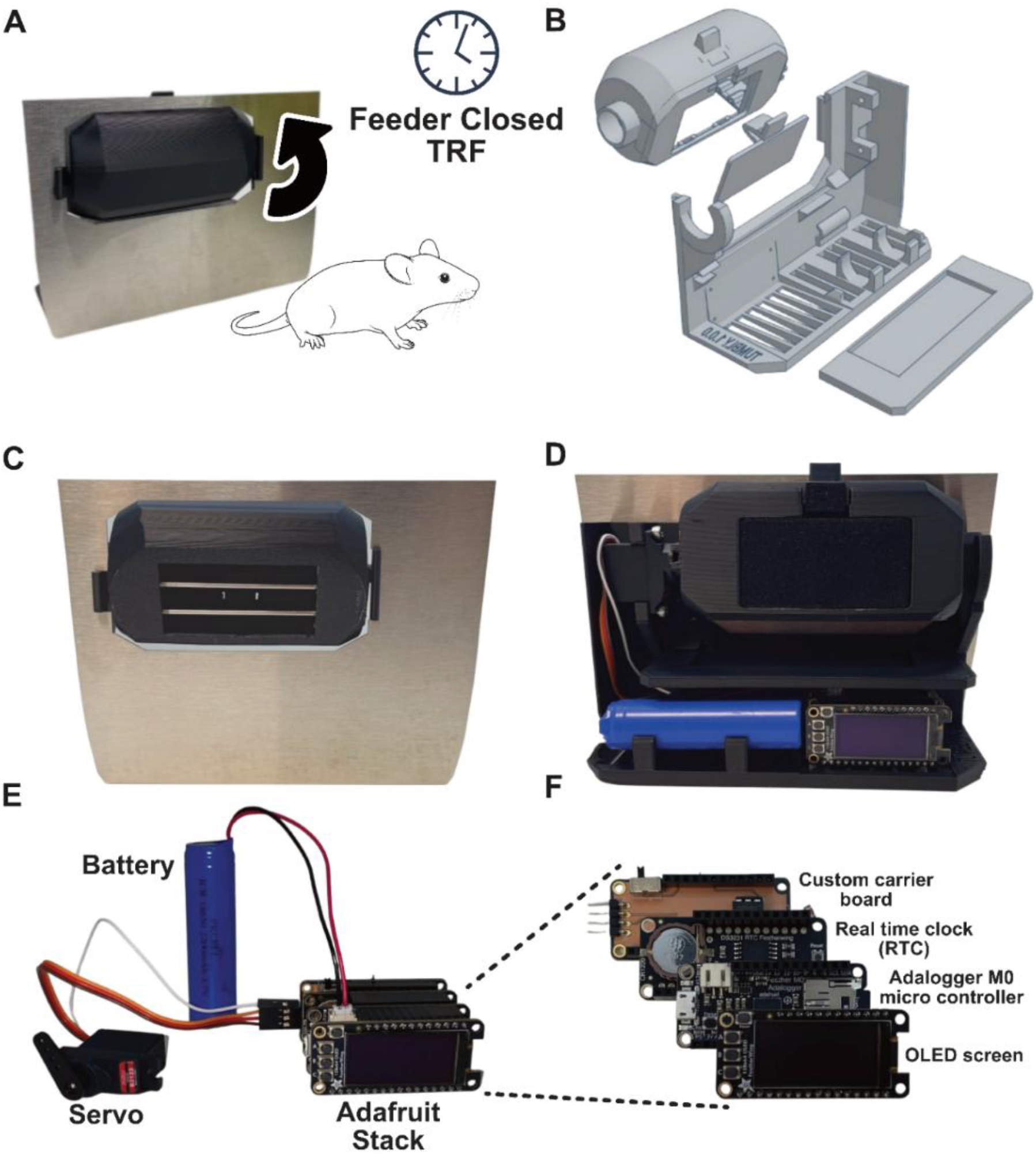
Overview of the Tumbly device. **A.** Schematic showing relative mouse size and the direction the food hopper opens. **B.** Exploded view of the 3D-printed parts: food hopper, hopper door, body, and dust cover. **C.** Front view with the hopper open. **D.** Back view. **E.** Electronics: servo, battery, and Feather stack. **F.** Boards in the Feather stack: custom carrier board, real-time clock, Feather M0, and OLED screen.

**Table 1.**
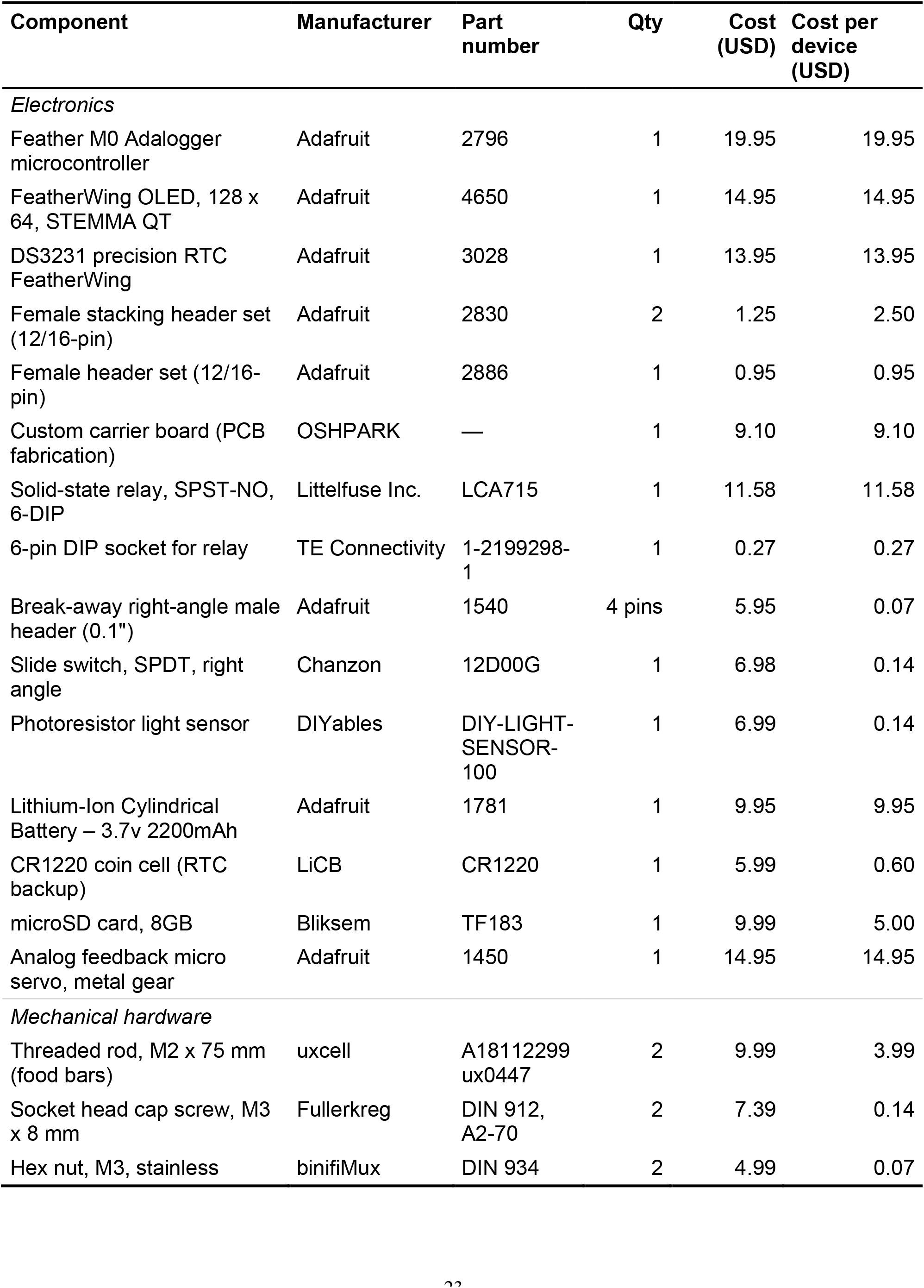
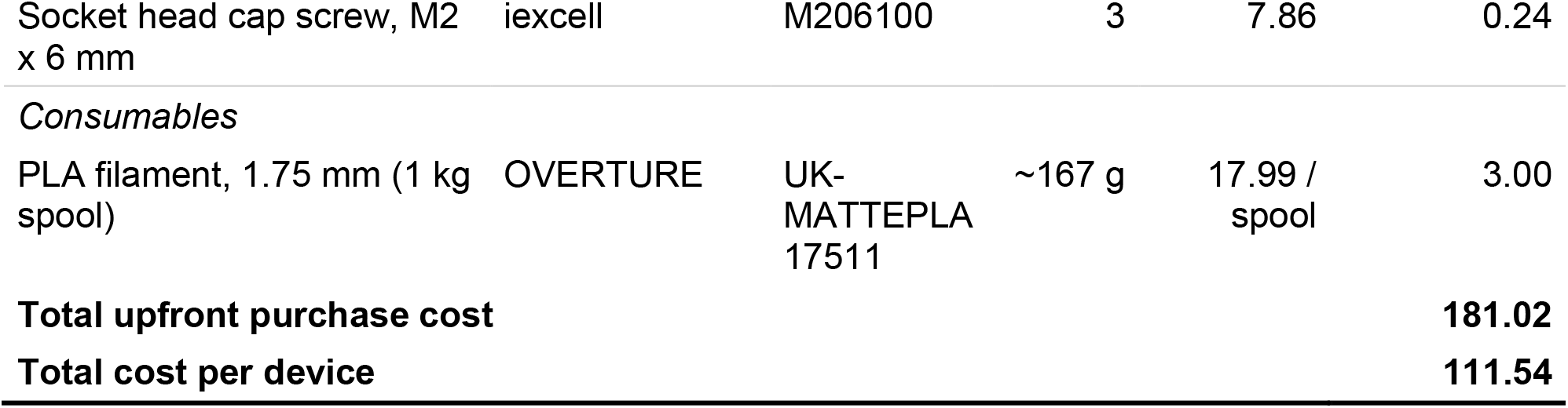
Bill of materials for a single Tumbly unit. All prices in USD as of July 2026, at single-unit purchase quantities from the listed vendors. Several components are sold only in multi-unit packs; unit cost for those items is the pack price divided by pack quantity, and cost per device is unit cost multiplied by the quantity required. Costs exclude 3D printing if outsourced, and laser cutting of the faceplate if selected in place of 3D printing.

### Electronics design

Tumbly electronics consist of three commercially sourced circuit boards and a custom-designed carrier printed circuit board (Fig. 1E,F). The stack begins with an Adafruit FeatherWing OLED screen on top, which contains three user-programmable buttons for controlling the user interface and device settings. Next is a DS3231 RTC FeatherWing for precise timekeeping. The final Adafruit module is the Feather M0 Adalogger microcontroller which runs the device firmware and includes a microSD card for device data logging. This Adafruit stack connects to a custom printed circuit board that contains a power switch, photodiode light sensor, LCA715 solid-state relay (SSR) chip for low-power servo control, and servo connector. The SSR cuts power to the servo while it is not in use, reducing power consumption and extending battery life.

### Firmware operation

User-controlled settings (device/mouse ID, task, open and close times, and open and close positions) are set using the three buttons and the OLED screen. Settings are written to a text file on the microSD card and reloaded at boot, so settings persist across power cycles or even battery removal. We programmed three tasks in the main code, which can be selected by the user: 1) Free Feeding, 2) TRF, and 3) Demo. Free Feeding always keeps the device open while logging light levels and food hopper position to the SD card every 30s, TRF opens and closes the food hopper based on the times set by the experimenter, again logging light levels and hopper position to the SD card every 30s, and the Demo program allows the user to test all device and servo functions.

Tumbly spends most of its operating life in a low-power “sleep” state to preserve battery life. Each wake cycle operates in a fixed order: read the light and battery level, execute hopper logic for the set task, check hopper position, log data, refresh the display, and sleep again for a configured interval (default 30 s). Data logged to the SD card every wake cycle includes datetime, device number, task, battery voltage, light level, hopper status, servo position, and potential errors. Every 30 min, the firmware also verifies that the hopper position is correct. If the servo cannot reach its target after moving (animal or bedding are obstructing movement), the firmware attempts several back-and-forth movements to disengage the obstruction. If these attempts fail, the device enters a fatal error state, the screen displays a persistent alert, further food hopper movement is suspended until an experimenter clears the obstruction and error with a button press.

### Battery runtime

Tumbly is powered by a single 3.7 V, 2200 mAh Li-ion cell (Adafruit, product 1781).

Using an Otii Arc power analyzer (Qoitech), we measured average current draws of approximately 60 mA while awake and 0.6 mA while sleeping. Battery runtime can be estimated from the duty-cycle-weighted average current:

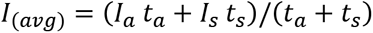

where *Iavg* is the duty-cycle-weighted average current, *Ia* and *Is* are the measured current draws during the awake and sleep states, respectively, and *ta* and *ts* are the durations of the awake and sleep states. The total runtime in days is approximately:

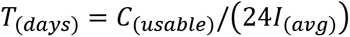

With the default 30 s sleep interval and an approximately 0.75 s active period per wake cycle, the average current is approximately 2.0 mA, corresponding to a theoretical runtime of approximately 45 days using the nominal 2200 mAh capacity. In six devices, empirical battery discharge corresponded to approximately 37 days of usable runtime, equivalent to approximately 82% of nominal battery capacity. This difference is consistent with the usable capacity being lower than the rated capacity because of battery discharge characteristics, voltage cutoff, and system losses. Therefore, the device checks battery voltage at each wake cycle and disables all processes below 3.5 V (equating to roughly 5-10% battery remaining) to reduce the risk of brownout-related filesystem corruption.

### Code accessibility

Tumbly runs on an open-source Arduino library (v1.5.0, released under GPL-3.0) available on GitHub and through the Arduino Library Manager (https://github.com/KravitzLabDevices/Tumbly). Other library dependencies are listed in the GitHub repository. Biological and device data can be found on a separate GitHub repository (https://github.com/MasBarr/Barrett2026) including Google Colab notebooks that run our analyses and create the figures used in this publication.

### Assembly

Building a Tumbly requires a 3D printer, a soldering iron, and M2 and M3 hex drivers.

We printed all parts on a Bambu H2D 3D printer in PLA filament, which took around 4.5 hours to print and used roughly 170 g of filament. The custom printed circuit board was ordered from OshPark (OshPark LLC) and components were soldered by hand in our laboratory. A full assembly video is available in the project repository (https://github.com/KravitzLabDevices/Tumbly/wiki). Tumbly was designed to be printed with PLA filament, which is not autoclavable. Between cohorts of animals, it is recommended that devices have external portions wiped down with standard lab cleaning agents (e.g. isopropyl alcohol) and tight corners blown with compressed air.

### Bill of materials and cost

The total cost to build one Tumbly unit is ∼$110 as of July 2026, at single-unit purchase quantities from the listed vendors (**Table 1**). This does not include the cost of outsourcing the 3D printing or laser cutting the faceplate, nor required tools to build Tumbly.

### Experimental animals

Device performance and biological validation were completed in two laboratories at separate institutions, using a total of 28 male mice (Lab A: n = 6, singly housed; Lab B: n = 22, group-housed in 7 cages). All procedures were approved by the Institutional Animal Care and Use Committee at Washington University in St. Louis and the Uniformed Services University of the Health Sciences (USUHS). All mice were housed under controlled temperature (24–25 °C) and humidity in their respective vivaria and had ad libitum access to water. Both cohorts were maintained on standard grain-based chow (LabDiet 5053 PicoLab Rodent Diet 20; 3.36 kcal/g, 4.5% of calories from fat).

### Behavioral Experiments

Lab A (Washington University in St. Louis) used six singly housed male mice on a human APOE4 knock-in background, approximately eight months of age at the start of the study, with no known pathology. In a within-subject design in which each mouse served as its own control, animals were first maintained under *ad libitum* feeding for seven days with the Tumbly device running the Free Feeding task, then transitioned to time-restricted feeding for seven more days with the food hopper open only during an eight-hour window (20:00–04:00; zeitgeber time (ZT) 14–22, where ZT0 is lights on). Mice were housed under a 12:12 light-dark cycle (18:00–06:00 dark). The middle day served as the transition day and was excluded from body weight and food intake analyses; activity recorded on that day was assigned to the ad libitum or TRF phase according to the time of the feeding switch. Each mouse was removed from its home cage and weighed individually on a laboratory scale once per day. Food consumption was recorded by weighing each Tumbly device daily and reported as grams per mouse per day. Locomotor activity was recorded continuously with passive infrared (PIR) sensors (Actimetrics Wireless Nodes, Model ACT-557-PIR) and analyzed with ClockLab.

Lab B (Uniformed Services University of the Health Sciences (USUHS)) used 22 male C57BL/6J mice (Jackson Labs, Bar Harbor, ME), group-housed at three to four animals per cage across seven cages, seven to eight weeks of age at the start of time-restricted feeding. In a between-group design, TRF (n = 12 mice, 4 cages) was compared to *ad libitum* feeding (n = 10 mice, 3 cages). All mice were acclimated to Tumbly devices under Free Feeding for two days, after which TRF cages were swapped to an eight-hour feeding window during the dark phase (17:00–01:00; ZT13–21) under a 12:12 light-dark cycle (16:00–04:00 dark). *Ad libitum* cages had the food hopper opened and were not powered, so device logs exist for the TRF units only. Food consumption was recorded as daily device weight per cage and reported as average grams per mouse per day. Individual body weight was recorded on days 0, 3, and 6.

### Statistical Analysis

Analyses were performed in Python using SciPy (v1.16.3) and statsmodels (v0.15.0) in Google Colab notebooks. Lab A used a within-subject, nonrandomized design in which each mouse experienced ad libitum feeding followed by TRF; the transition day was excluded from body weight and food intake analyses. Body weight (last day of each phase) and mean daily food intake were compared with paired t-tests. Light- and dark-phase activity counts were summed per day, averaged within each phase, and analyzed with a two-way repeated-measures ANOVA (feeding condition × light/dark period). Lab B used a between-group design in which cages were randomly assigned to TRF or ad libitum feeding. Because mice within a cage are not independent, cage was the unit of analysis: last-day body weight (cage mean) and mean daily food intake per mouse (cage intake divided by mice per cage) were compared with unpaired t-tests. Given the small sample sizes, normality was assumed and parametric tests were used.

Sample sizes were not determined by power analysis and reflect the number of devices and animals available for validation. Significance was set at α = 0.05 (two-tailed). Data are presented as mean ± SEM unless otherwise noted; hopper timing delays are presented as mean ± SD. Effect sizes and 95% confidence intervals for all tests are reported in Table 2.

**Table 2.** Statistical table.

|  | Data structure | Type of test | Effect size and 95% CI |
| --- | --- | --- | --- |
| **a** | Normality assumed, n = 6<br>mice | Paired t-test | 0.38 g [−0.63, 1.40] |
| **b** | Normality assumed, n = 6<br>mice | Paired t-test | 0.26 g/day [−0.23, 0.74] |
| **c** | Normality assumed, n = 6<br>mice | Two-way RM ANOVA,<br>condition | 1,177 counts [444, 1,911];<br>$\eta p^2 = 0.77$ |
| **d** | Normality assumed, n = 6<br>mice | Two-way RM ANOVA,<br>period | 3,260 counts [2,408, 4,112];<br>$\eta p^2 = 0.95$ |
| **e** | Normality assumed, n = 6<br>mice | Two-way RM ANOVA,<br>interaction | −52 counts [−1,761, 1,657];<br>$\eta p^2 < 0.01$ |
| **f** | Normality assumed, n = 3<br>vs. 4 cages | Unpaired t-test | 0.30 g [−1.16, 1.76] |
| **g** | Normality assumed, n = 3<br>vs. 4 cages | Unpaired t-test | 0.18 g/mouse/day [−0.25,<br>0.61] |

Effect sizes are mean differences (ad libitum – TRF) with 95% confidence intervals; for d, the period contrast is dark – light. Letters correspond to superscripts in the Results text.

## AI statement

Claude (Anthropic) was used in the analysis of the data presented. All analysis was verified by the authors and is available on GitHub.

## Results

Device performance was assessed in two different labs (6 units tracking 6 mice in Lab A, 7 units tracking 22 mice in Lab B). In Lab A (Fig. 2A), body weight did not differ significantly between ad libitum feeding and TRF phases (33.1 ± 0.6 g vs. 32.7 ± 0.8 g, paired t(5) = 0.970, p = 0.377^a^; Fig. 2C), indicating that mice obtained adequate food in both ad libitum and TRF phases (Fig. 2B). Daily food intake dropped on the first day of restriction and recovered over the following days (Fig. 2D), averaging 3.76 ± 0.09 g/day under ad libitum feeding and 3.50 ± 0.16 g/day under TRF (paired t(5) = 1.368, p = 0.230^b^; Fig. 2E). Home-cage activity was monitored with PIR sensors and remained concentrated in the dark phase under both conditions (Fig. 2F). Total daily activity was lower under TRF in both the light and dark phase (light phase: 4284 ± 1061 vs. 3133 ± 772 counts; dark phase: 7570 ± 1224 vs. 6366 ± 1128 counts; 2 × 2 repeated-measures ANOVA: main effect of feeding condition F(1, 5) = 17.03, p = 0.009^c^, main effect of period F(1, 5) = 96.67, p = 0.0002^d^, condition × period interaction F(1, 5) = 0.01, p = 0.940^e^; Fig. 2G). Reduced activity under TRF has also been reported by other groups (Whittaker et al., 2023), suggesting this is a common effect of TRF rather than an artifact of the device.

**Figure 2.**
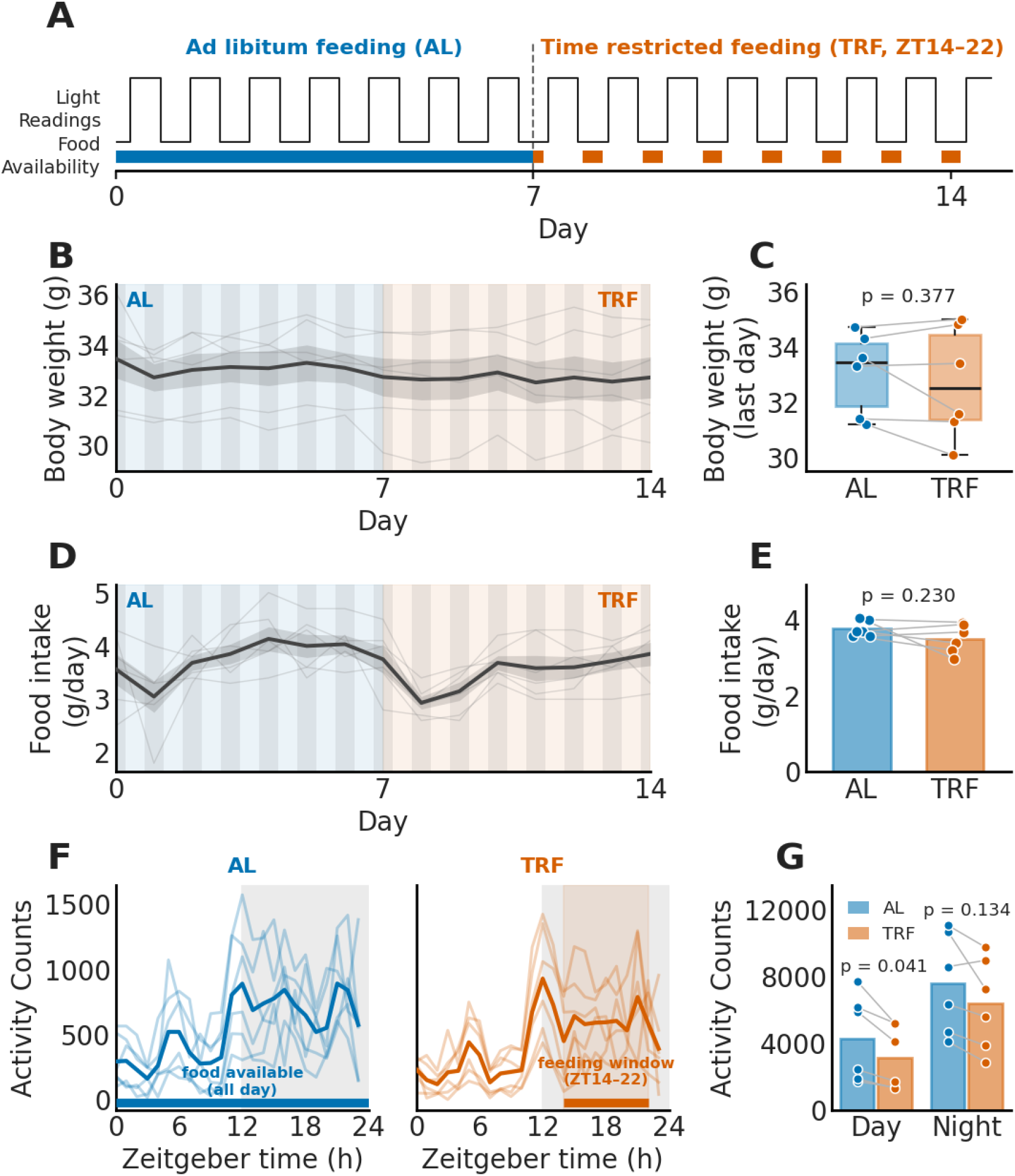
Validation in Lab A (singly housed, within-subject; n = 6 males). **A.** Experimental timeline and TRF schedule (ZT14-22, 20:00–04:00). **B.** Line plot showing daily body weight over 14 days, mice were swapped from *ad libitum* (AL) to TRF on day 7. **C.** Box and whisker plot showing average and individual body weight for the last day of AL and last day of TRF. **D.** Line plot showing daily food intake. **E.** Bar plot showing average and individual food intake in each phase. **F.** Line plots showing average hourly activity AL (left) and TRF (right) phases. **G.** Bar plot showing average and individual activity counts during the light and dark phases under AL and TRF. Thin lines and points show individual mice; thick lines and bars show means; shaded bands show SEM. Grey vertical bands in B and D mark the dark phase within each day, and the blue and orange background tints mark the AL and TRF periods, respectively. Statistics in C and E: paired t-tests; G: two-way repeated-measures ANOVA (feeding condition × light/dark period).

In Lab B (Fig. 3A), the TRF and ad libitum groups began at equivalent body weight (ad libitum 24.45 ± 0.41 g, TRF 24.23 ± 0.34 g) and ended at 24.74 ± 0.36 g and 24.44 ± 0.41 g (ad libitum +0.29 ± 0.25 g, TRF +0.20 ± 0.16 g; unpaired t(5) = 0.533, p = 0.617^f^; Fig. 3B,C). Food intake in the TRF group was maintained on the first day of restriction (3.62 ± 0.13 g/mouse/day), dropped on the second day (2.42 ± 0.26 g/mouse/day), and recovered to control levels by day 4 (Fig. 3D). Averaged over the 7 days, intake was 3.35 ± 0.11 g/mouse/day under TRF and 3.53 ± 0.12 g/mouse/day under ad libitum feeding (unpaired t(5) = 1.079, p = 0.330^g^, n = 4 and 3 cages; Fig. 3E). Home-cage activity levels were not measured in Lab B.

**Figure 3.**
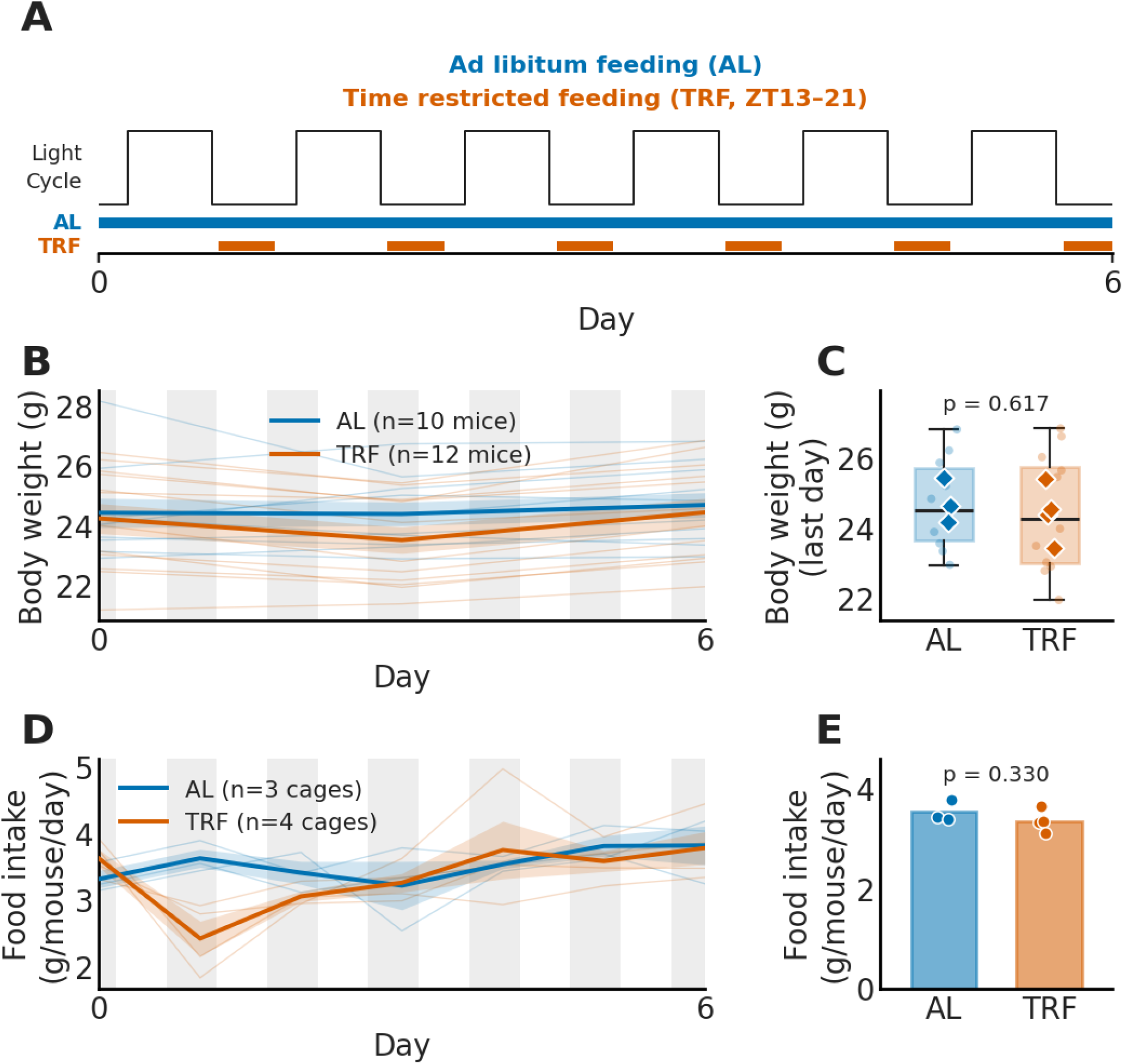
Validation in Lab B (group-housed, between-group; AL: n = 10 males in 3 cages, TRF: n = 12 males in 4 cages). **A.** Experimental timeline and TRF schedule (ZT13-21, 17:00–01:00). **B.** Line plot showing body weight over 7 days. **C.** Box-and-whisker plot of last-day body weight; faint points, individual mice; diamonds, cage means (the analysis unit). Statistics on cage means (n = 4 TRF vs 3 AL). **D.** Line plot showing daily food intake per mouse, calculated as cage intake divided by the number of mice. **E.** Bar plot showing average food intake, each point is one cage (cage intake divided by the number of mice). Thin lines show individual mice (B) or cages (D); thick lines and bars show means; shaded bands show SEM. Grey vertical bands in B and D mark the dark phase within each day. Statistics in C and E: unpaired t-tests.

We quantified potential device errors by examining data log files for food hopper movement times and errors at the conclusion of the two experiments. Scheduled hopper transitions occurred shortly after their programmed times, with a mean delay of 17.9 ± 9.1 s for openings and 18.9 ± 9.9 s for closings. 57 position errors were detected across all experiments, and all were detected and resolved automatically by Tumbly moving the food hopper to achieve the calibrated position. This resulted in an average of ∼1 correction per 2 days per device.

Importantly, no errors stopped device functioning, totaling >1000 hours of operation without a fatal device error.

## Discussion

Here, we demonstrate Tumbly, a new open-source home cage device for TRF experiments. Tumbly was adapted from the Tumble Feeder, an open-source home-cage device that we previously developed to control access to high-fat diet in response to nose-pokes (Barrett et al., 2025). We improved the design of Tumbly to be easier to build, and optimized for TRF.

Commercial systems such as BioDAQ and ClockLab Feeder Control can control and measure intake at the bout or pellet level, logging data with dedicated software. However, these require specific caging and wired connections, precluding their use on typical vivarium housing racks. Several open-source systems have also been developed to perform TRF (Greenwell et al., 2019; Sahasrabudhe et al., 2022; Farré et al., 2024; Lê et al., 2025), each with distinct strengths. Tumbly complements these devices with a small and wire-free design, and smart features such as closed-loop position verification and automatic error recovery (which was used to adjust the position 57 times in the current studies). Tumbly is also relatively affordable (∼$110/device), which makes it ideal for labs with smaller budgets, or labs that want to perform high throughput TRF studies on vivarium racks. Finally, Tumbly is open-source and includes onboard STEMMA connectors that can support additional environmental sensors, such as CO2, temperature, or humidity sensors, allowing the platform to be adapted for more complex experimental applications.

While we highlight the strengths of Tumbly in this paper, there are also limitations. First, because Tumbly is battery powered, experiments are ultimately limited by battery life and require charging or battery replacement every ∼5 weeks for long studies. Second, Tumbly was designed to control food access and not precisely control meal size or total caloric intake. As a result, Tumbly is not suitable for experiments requiring delivery of exact quantities of food at specific times. Related to this point, Tumbly cannot weigh the amount of food removed.

Therefore, food intake must be estimated by weighing the Tumbly device itself to determine how much food was removed. This unfortunately cannot distinguish food eaten from food spilled or hoarded by the mouse. The dust cover beneath the hopper partially addresses this by catching small fragments of diet which can be accounted for during weighing. But ultimately, Tumbly is not suited to precise control or quantification of caloric intake. Third, to keep the device simple and low cost, we also did not include touch or beam-break sensing at the food hopper, so the device cannot quantify feeding bouts or patterns as the “Tumble Feeder” it was based on did (Barrett et al., 2025). Finally, because data are stored locally on a microSD card, Tumbly does not currently provide real-time remote monitoring or centralized data collection, meaning device errors can only be discovered during routine checks rather than immediately. Although we did not experience any fatal errors in our validation tests, it is critical for researchers to check Tumbly daily to ensure that an error has not occurred that would deprive the mice of food. We hope at least some of these limitations can be overcome in future iterations of Tumbly, which build on its open-source design.

## Acknowledgements

We thank members of the Kravitz and Creed labs for their feedback and advice throughout this project.

---

**Movie 1.** Tumbly opening and closing on its programmed schedule in the home cage with a mouse present.

