## Supplementary figures and images for "Tumbly: An Open-Source Home-Cage Device for Time-Restricted Feeding in Mice"

### Movie 1

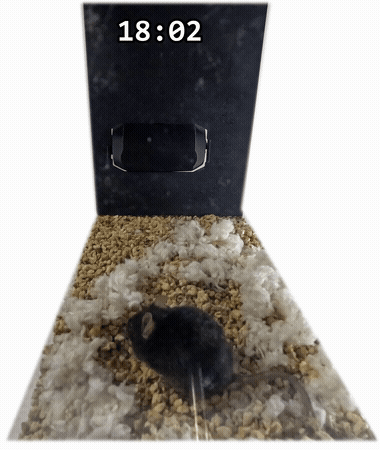
